# Intraspecific Variation in Gulf Killifish (*Fundulus grandis*) Avoidance of Contaminants

**DOI:** 10.1101/2025.08.04.668507

**Authors:** Charles W. Martin, Ashley M. McDonald, Kathryn A. O’Shaughnessy

## Abstract

Understanding the behavioral response of estuarine fish populations to environmental contaminants is critical for assessing the ecological consequences of disturbances. Here, we examined population-level variation in avoidance behavior to petroleum-contaminated sediments in the Gulf killifish (*Fundulus grandis*), a sentinel species in northern Gulf of Mexico marshes. Adult fish were collected from four geographically distinct estuaries—Terrebonne Bay, Louisiana; Mississippi Sound, Alabama; Pensacola Bay, Florida; and Cedar Key, Florida. Choice experiments tested fish preference to either uncontaminated sediments or sediments contaminated with fresh or weathered oil. Results found significant differences among populations with individuals from western Gulf sites (Louisiana and Alabama) exhibiting strong avoidance of fresh oil, while those from eastern sites (Florida) showed limited or no avoidance. No populations avoided weathered oil. These findings suggest that sensitivity to oil contamination varies regionally, potentially reflecting local exposure histories or environmental adaptation. The diminished response to weathered oil across all populations highlights a reduced ability to detect residual contamination and indicates that these fish may use more volatile compounds that precipitate or degrade faster as a cue. This study underscores the importance of incorporating behavioral and population-level variability into impact assessments and emphasizes the need for regionally tailored conservation strategies.

## Introduction

Estuarine ecosystems provide numerous ecosystem services, ranging from storm protection (Barbier, 2020) to nutrient reduction (Seitzinger, 1988) and enhanced production of commercially and recreationally important species (Beck et al., 2001). Salt marshes are widespread and vital estuarine habitats that facilitate energy transfer to adjacent coastal waters. Due to the highly dynamic conditions of estuaries, these systems often depend on a limited number of resident taxa (Day et al., 1989; Whitfield, 1994). The estuarine resident Gulf killifish (*Fundulus grandis*) is a mid-trophic level species essential for maintaining trophic connectivity and ecosystem resilience (Subrahmanyam and Drake, 1975; Rozas and LaSalle, 1990; Costanza et al., 1993; Tett et al., 2013). *Fundulus grandis* is a common and ecologically important resident of northern Gulf of Mexico salt marshes and has become a sentinel species for assessing environmental disturbance due to its abundance (Able et al., 2015), high site fidelity (Nelson et al., 2014; Vastano et al., 2017; Jensen et al., 2019), and central role in food webs (McCann et al., 2017; Oken et al., 2023). Its hardiness, ease of capture, and ability to thrive in captivity also make it an ideal species for laboratory studies of environmental stressors (Whitehead et al., 2012; Dubansky et al., 2013, Dubansky et al., 2017).

Populations of estuarine fishes often experience localized environmental conditions and stressors that can shape physiological tolerance and behavioral traits, including responses to contaminants.

Previous studies have shown that population-level differences in sensitivity to environmental toxins can arise from both genetic differentiation and phenotypic plasticity (Whitehead et al., 2012; Schlenker et al., 2019). For *F. grandis*, such variability could result in differential susceptibility to hydrocarbon exposure, depending on historical oiling, contaminant loads, or other co-occurring stressors such as hypoxia and salinity fluctuations. As a result, individuals from different populations may vary in their capacity to detect and avoid contaminated sediments, with potential consequences for habitat use, exposure risk, and resilience to oiling events (Fodrie et al., 2014). Understanding this variability is crucial for evaluating how widespread disturbances like oil spills affect estuarine species at scales relevant to conservation and management.

The 2010 *Deepwater Horizon* (DwH) oil spill released approximately 4.9 million barrels of crude oil into the northern Gulf, impacting over 1,700 km of shoreline, predominantly saltmarshes (McNutt et al., 2012; Michel et al., 2013; Nixon et al. 2016). Initial predictions anticipated severe ecological disruption, particularly to nearshore fish communities. However, subsequent field surveys often found minimal to no long-term population-level impacts on coastal fishes (Fodrie and Heck, 2011; Moody et al., 2013; Fodrie et al., 2014; Able et al., 2015; Schaeffer et al., 2016). This apparent resilience has been attributed to several factors, including behavioral avoidance of contaminated habitats, patchy oil distribution, and rapid recolonization from unaffected areas (Fodrie et al., 2014; Martin, 2017; Martin et al., 2020a). Laboratory experiments, however, have consistently documented sublethal physiological and behavioral effects of oil exposure on estuarine fishes, including reduced foraging efficiency, impaired locomotion, and altered olfactory responses (Clairaux et al., 2004; Whitehead et al., 2012; Dubansky et al., 2013; McDonald et al., 2022). Specifically, *F. grandis* demonstrates strong avoidance of fresh oil-contaminated sediments, but this response is diminished with weathered oil or after prior exposure to oil, suggesting potential impairment of sensory detection mechanisms (Martin, 2017; Martin et al., 2020b). These behavioral alterations may impact habitat selection, foraging success, and ultimately, fitness.

Despite the growing body of evidence for behavioral responses to oil exposure, little is known about intraspecific variation in these responses across geographically or environmentally distinct populations. Given the extensive spatial heterogeneity in oiling during the DwH spill and potential for localized adaptation or acclimatization, it is critical to examine whether populations differ in their ability to detect and avoid oil-contaminated habitats. Such variation could have significant implications for population resilience and the persistence of ecological functions in impacted areas. In this study, we experimentally test the behavioral responses of multiple *F. grandis* populations to varying concentrations of oil-contaminated sediments. We hypothesize that (1) fish will exhibit avoidance of fresh oil-contaminated sediments relative to uncontaminated controls or weathered oil-contaminated sediments, and (2) the strength of avoidance behavior will vary among populations, potentially reflecting differences in previous exposure history or local environmental adaptation.

## Methods

### Fish Collection

Adult (>65mm total length) Gulf killifish (*F. grandis*) were collected from four geographically distinct estuarine locations along the northern Gulf of Mexico: Terrebonne Bay, Louisiana (TB); Mississippi Sound, Alabama (MS); Pensacola Bay, Florida (PB); and Cedar Key, Florida (CK) (Figure 1A). At each site, fish were captured using baited minnow traps placed along the marsh edge in shallow intertidal zones. After capture, fish were transported to the laboratory and housed in aerated, flow-through holding systems maintained at salinity levels consistent with collection conditions. To minimize potential artifacts associated with capture stress and handling, all fish were held for a minimum of one month prior to behavioral testing and acclimated to 8 psu, a common salinity found in Gulf estuaries. During this acclimation period, fish were fed daily and monitored for health (lesions, injuries, etc.) and abnormal behavior. Only healthy fish were used in trials and each fish was used once during experiments. Collection and use of fish in experiments was conducted with IACUC approval #201710044.

**Figure 1.**
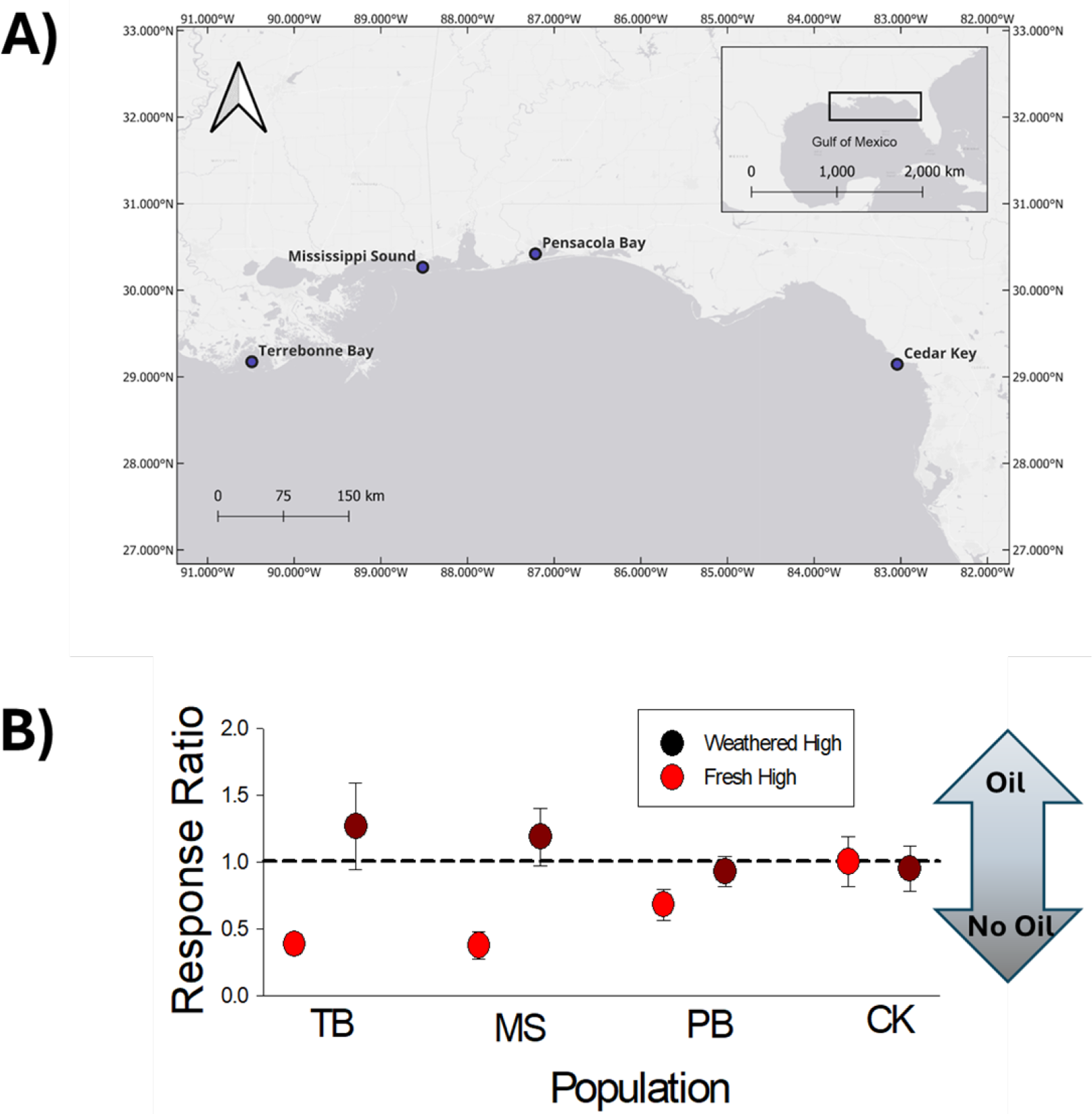
Gulf Killifish (*Fundulus grandis*) was collected from Terrebonne Bay, Louisiana (TB); Mississippi Sound, Alabama (MS); Pensacola Bay, Florida (PB); and Cedar Key, Florida (CK) (A). Fish demonstrated variability in their response ratio (B), with the dashed line (1.0) indicating no preference among weathered or fresh oil choices and avoidance of oil <1.0.

### Experimental Trials

Behavioral avoidance trials followed the protocol described in Martin (2017). Trials were conducted in 38-liter aquaria divided into two equal zones: one side containing clean sediment and the other side containing sediment mixed with oil. Approximately 3 L of sediment (1.5 L per side) was added to each tank and leveled to a uniform depth of ∼18 cm. For choices involving oil, the oil was thoroughly mixed with the sediment prior to placement and then covered with ∼2 cm of clean sediment to simulate shallow burial and prevent oil from surfacing during trials. Two oil treatments were tested: fresh oil and weathered oil, each at high concentrations (40 mL oil L^−1^ sediment), which represents the upper range of contamination reported in coastal habitats following the *Deepwater Horizon* spill (Turner et al., 2014) and known to induce behavioral response (Martin, 2017).

Weathered oil was produced by aerating fresh crude in a fume hood to achieve 40% mass loss, mimicking conditions typical of oil that had undergone environmental weathering prior to marsh deposition (specific sediment and aromatic concentrations are reported in Martin, 2017).

For each trial, a single fish was introduced to the center of the tank and allowed to acclimate for five minutes. Following acclimation, fish behavior was recorded using a GoPro digital camera for a 10-minute observation period. Fish position was recorded every 30 seconds, resulting in a total of 20 observations per trial. Occupancy was calculated as the proportion of observations in which the fish was located on each side of the tank, and then the ratio of oiled to unoiled proportions compared among populations and oil types as the response variable.

Side assignment (left or right) for oiled sediment was randomized across trials to control for any side bias. Trials in which fish did not swim or remained motionless for the duration were excluded from analysis. Behavioral responses were then compared using a two way Analysis of Variance (ANOVA) among treatments of population (fish from the four collection sites) and oil treatments (fresh vs. weathered), as well as the interaction, after meeting assumptions of the test (homogeneity of variance and normality). When normality failed, a Kruskal-Wallis test was used. Each unique population * oil treatment was replicated 12 times and no single fish was used in multiple trials.

## Results

A two-way ANOVA revealed a significant interaction between population and oil treatment on response ratio (F_3,88_ = 3.38, p = 0.0217), indicating that the behavioral response to oil depended on both population origin and oil condition (fresh vs. weathered) (Figure 1B). Significant behavioral responses to oil-contaminated sediments were detected between oil treatments (F_1,87_ = 3.38, p = 0.0217), with fresh oil triggering strong avoidance responses while fish in weathered oil did not have preferences. The population factor did not significantly vary among *F. grandis* populations (F_3,87_ = 0.582, p = 0.629) presumably due to the strong and variable effect of fresh versus weathered oil on behavior. When testing just fresh oil, strong differences among populations existed (Kruskal-Wallis; H = 11.48, p = 0.009).

Fish from Mississippi Sound, Alabama (MS) and Terrebonne Bay, Louisiana (TB) showed the strongest avoidance of fresh oil, with mean response ratios well below 1.0 (MS: 0.38 ± 0.10 SE; TB: 0.42 ± 0.09 SE), indicating that individuals from these populations spent proportionally less time over oiled sediments. In contrast, fish from Pensacola Bay, Florida (PB) and Cedar Key, Florida (CK) did not exhibit significant avoidance, with mean response ratios near or slightly above 1.0 (PB: 0.68 ± 0.12 SE; CK: 1.00 ± 0.18 SE). No population exhibited significant avoidance of weathered oil. Across all locations, fish displayed approximately equal preference for oiled and clean sediments under weathered conditions, with mean response ratios near 1.0 (MS: 1.19 ± 0.21 SE; TB: 0.93 ± 0.15 SE; PB: 0.71 ± 0.09 SE; CK: 0.95 ± 0.17 SE), suggesting that behavioral avoidance is specific to fresh oil exposure and may be diminished or absent when fish encounter weathered hydrocarbons.

## Discussion

Our results demonstrate population-specific sensitivity to oil contamination, with western Gulf populations (MS, TB) displaying stronger avoidance responses than eastern populations (PB, CK). These findings suggest potential regional variation in behavioral plasticity, prior exposure history, or adaptive responses to oil contamination. This geographic variability in behavioral sensitivity suggests that population-specific traits, potentially influenced by multiple generations and natural selection to local environmental histories or potentially differential exposure to contaminants, may influence ecological resilience following oil spills. Additionally, the consistent lack of avoidance across all populations to weathered oil highlights a reduction in perceived risk or detectability of hydrocarbons after environmental degradation, with important implications for exposure risk in marshes where weathered oil persists long after acute spill events.

The observed west-to-east gradient in avoidance responses may reflect spatial differences in prior exposure to oil, especially given that marshes in Louisiana, Mississippi, and Alabama received disproportionately higher oil loads during the DwH spill (Michel et al., 2013). Outside of major spills such as the Exxon Valdez (Peterson et al., 2003), Ixtoc (Soto et al., 2014), Arthur Kill (Burger, 1994), and Falmouth Harbor (Reddy et al., 2002), smaller spills are frequent -especially in the northern Gulf of Mexico. The United States Coast Guard National Response Center has over 20,000 spills reported annually (Martin et al., 2023) and in Louisiana alone the Louisiana Oil Spill Coordinator’s Office (LOSCO) responded to 44 spills in 2024 (https://data.losco.org). Previous exposure to hydrocarbons has been shown to alter olfactory detection, risk assessment, and overall behavioral responses in fishes (Martin et al., 2020b; Schlenker et al., 2019). Thus, fish from heavily oiled regions or areas with frequent leaks/spills, which likely experience sublethal exposures, may have retained or developed stronger avoidance behavior as an adaptive or conditioned response.

Alternatively, these differences could arise from local adaptation or phenotypic expression driven by unrelated environmental variability (e.g., salinity, hypoxia, or anthropogenic pollutants; Hamilton et al., 2016), which may influence the development and expression of avoidance behavior across populations.

Our findings align with earlier work demonstrating that *F. grandis* avoid heavily oiled sediments in laboratory trials (Martin, 2017; Martin et al. 2020b), but add a new dimension by revealing that such responses are not uniform across the species’ range. This variation has important ecological implications: if certain populations are less likely to detect and avoid contamination, they may be more susceptible to sublethal effects on foraging, reproduction, and predator avoidance (Rice et al., 1987; Khursigara et al., 2021), thereby enhancing population- or community-level resilience.

Furthermore, populations with reduced avoidance responses may be at greater risk of prolonged exposure to harmful compounds, even in weathered oil conditions that do not elicit immediate behavioral aversion but can still impair physiological function (Whitehead et al., 2012).

The observed lack of avoidance to weathered oil may yet have significant implications for *F. grandis* populations, given that weathering increases the proportion of high molecular weight polycyclic aromatic hydrocarbons (PAHs) in crude oil (Di Toro et al., 2007). Most studies investigating toxicity of weathered versus fresh oil have focused on early life stages, finding weathered oil more harmful to fish embryos and larvae than fresh oil (Incardona et al., 2014; Esbaugh et al., 2016; O’Shaughnessy et al., 2018); but these results may still be relevant to later life stages. Although they did not compare to fresh oil, Lee et al. (2018) found significant immune dysfunction of rockfish (*Sebastes schlegeli*) after oral administration of weathered Iranian heavy crude oil. It might be that adults, however, are less susceptible to these effects or have escaped effects completely, possibly due to developmental stage-specific sensitivity of weathered oil.

Several caveats must be considered when interpreting these results. First, the laboratory conditions used in the avoidance trials simplify the complex structural, chemical, and ecological dynamics of natural marsh environments. Moreover, we acknowledge the absence of vegetation, conspecifics, and flow may limit the ecological realism of the tests. The use of laboratory experiments offers a tradeoff between increased control of environmental conditions compared to field observations/manipulations, which are often plagued by the difficulty in teasing apart many co-occurring factors, some of which may be unmeasured (Diamond 1986). In a review of studies of DwH effects, laboratory/mesocosm experiments contributed 38% of all studies (Martin et al., 2023). Second, while we held fish in captivity for a month to minimize handling artifacts, laboratory acclimation may still influence behavior in subtle ways. Third, the response ratio metric is based on spatial occupancy and does not incorporate more nuanced behavioral cues such as swimming speed, gill ventilation, or exploratory behavior, which could provide additional insight into avoidance dynamics. Fourth, sample sizes (n=12 for each unique treatment) for some treatment combinations, while adequate for detecting strong patterns, may have limited our ability to detect more subtle interactions. Finally, it is also important to note that these trials represent short-term, acute exposures; longer grow-out periods following exposure, as used in other studies (e.g., O’Shaughnessy et al., 2018; Vignet et al., 2019), may reveal delayed or latent effects that would otherwise go undetected in short-duration tests.

Overall, this study underscores the importance of incorporating population-level variation into assessments of oil spill impacts. Behavioral traits such as habitat avoidance may buffer some populations against acute contamination, while others may face higher exposure risk due to diminished sensitivity. Future studies should investigate the mechanistic basis of these differences, whether genetic, developmental, or experiential, and explore how they interact with other environmental stressors. Understanding this variability will be essential for predicting estuarine fish responses to future disturbances and informing site-specific conservation or restoration strategies.

## Acknowledgments

We thank the members of the Coastal Waters Consortium for their support in this work. Moreover, we thank Placid Refining Company for their provision of Louisiana Sweet Crude used in mesocosm oiling efforts.

## Data Availability Statement

Studies used in the analysis are publicly available through the Gulf of Mexico Research Initiative Information and Data Cooperative (GRIIDC) at https://data.gulfresearchinitiative.org.

